# The ins and outs of membrane bending by intrinsically disordered proteins

**DOI:** 10.1101/2022.12.17.520884

**Authors:** Feng Yuan, Christopher T. Lee, Arjun Sangani, Justin R. Houser, Liping Wang, Eileen M. Lafer, Padmini Rangamani, Jeanne C. Stachowiak

## Abstract

Membrane curvature is essential to diverse cellular functions. While classically attributed to structured domains, recent work illustrates that intrinsically disordered proteins are also potent drivers of membrane bending. Specifically, repulsive interactions among disordered domains drive convex bending, while attractive interactions, which lead to liquid-like condensates, drive concave bending. How might disordered domains that contain both repulsive and attractive domains impact curvature? Here we examine chimeras that combine attractive and repulsive interactions. When the attractive domain was closer to the membrane, its condensation amplified steric pressure among repulsive domains, leading to convex curvature. In contrast, when the repulsive domain was closer to the membrane, attractive interactions dominated, resulting in concave curvature. Further, a transition from convex to concave curvature occurred with increasing ionic strength, which reduced repulsion while enhancing condensation. In agreement with a simple mechanical model, these results illustrate a set of design rules for membrane bending by disordered proteins.

## Introduction

Highly curved membrane surfaces are found throughout the cell and play a role in a myriad of cellular processes, from endocytosis and exocytosis, to budding of enveloped viruses, protein recycling during autophagy, and the structure and maintenance of all organelles(*1-3*). Protein-lipid interactions are known to drive membrane curvature through several established and emerging mechanisms. The first mechanisms of curvature generation to be characterized relied upon proteins with specific structural features. For example, insertion of a wedge-like amphipathic helix into the membrane surface increases the area of one membrane leaflet relative to the other, causing the membrane to bend toward the protein layer, such that convex membrane buds and tubules “coated” by proteins are created(*4*). In a second mechanism, proteins that bind to membrane surfaces using inherently curved surfaces, such as BAR (Bin/Amphiphysin/RVS) domains, can drive the membrane to conform to their curvature. Interestingly, these scaffolds can have either convex or concave surfaces, enabling them to produce either protein-coated or protein-lined membrane buds and tubules, respectively(*5*).

More recent work has demonstrated that specific structural motifs, such as amphipathic helices and BAR domains, are not the only means of generating membrane curvature. In particular, several reports have demonstrated that proteins without a well-defined secondary structure - intrinsically disordered proteins - are also capable of shaping membrane surfaces(*6, 7*). Several proteins involved in endocytosis, including AP180 and Epsin1, contain intrinsically disordered domains with substantial molecular weight (400-500 amino acids) and high net charge. When these domains become crowded on membrane surfaces, steric and electrostatic repulsion between them drives the membrane to bend toward the protein layer, such that the area available per protein domain is increased. This process leads to convex, protein-coated membrane buds and tubules(*7, 8*). Similarly, crowding among glycosylated proteins on the plasma membrane surface is thought to drive assembly of tube-like cellular protrusions(*9*).

In contrast, many disordered domains have recently been found to attract one another through a network of weak interactions, leading to condensation of a protein liquid phase(*10, 11*). When disordered domains with these attractive interactions encounter one another on membrane surfaces, they seek to maximize contact with one another, generating a compressive stress at the membrane surface(*12*). This stress bends the membrane away from the protein layer such that the area per protein on the membrane surfaces is decreased, resulting in concave, protein-lined membrane buds and tubules.

These observations collectively suggest that the differential stresses induced by a layer of disordered proteins on the membrane surface can be tuned to control the directionality and magnitude of membrane bending. Importantly, disordered domains that are repulsive or non-interacting and disordered domains that attract one another, often exist within the same protein, and have been characterized as “stickers” and “spacers”, respectively, by computational modeling efforts(*13*). Therefore, to predict the overall impact of an intrinsically disordered protein on membrane curvature, we must understand how repulsive and attractive domains work together to apply bending stresses to the membrane surface.

Toward this goal, here we examine a series of disordered protein chimeras, which combine protein domains previously shown to drive either convex or concave membrane curvature. Using these chimeras, we demonstrate that disordered protein layers with opposite curvature preferences can either work together to amplify curvature or can oppose one another to create context-dependent control of membrane shape. In agreement with a simple mechanical model, this work outlines a set of design rules that can be used to understand the impact of disordered proteins on membrane curvature.

## Results

### Attractive domains cluster repulsive domains at membrane surfaces, amplifying convex membrane bending

Here we examine a series of recombinant protein chimeras which link disordered protein domains that predominantly repel one another with disordered domains that predominantly attract one another. For the repulsive domain, we have chosen the C-terminal domain of the endocytic adaptor protein, AP180. Previous work has demonstrated that this domain generates repulsive interactions at membrane surfaces through a combination of steric and electrostatic effects(*14*). We used amino acids 328-518 of AP180, approximately the first third of the C-terminal domain, which has a net negative charge of -21(*14*). We will refer to this domain henceforth as the “short” version of AP180, or AP180S. For the attractive domain, we chose the low complexity domain of fused in sarcoma (FUSLC), residues 1-163. FUSLC is known to undergo liquid-liquid phase separation (LLPS) both in solution(*15*) and when recruited to membrane surfaces(*12*). FUSLC domains attract one another through a combination of p-p and dipole-dipole interactions among amino acid side chains(*10*).

The first chimera we examined consisted of an N-terminal histidine tag, for attachment to DGS-NTA-Ni lipids, followed by the FUSLC and AP180S domains, FUSLC-AP180S (Figure 1a left panel). When this protein attaches to the membrane surface using its histidine tag, the FUSLC domain is closer to the membrane relative to AP180S (Figure 1a right panel). The individual domains, his-AP180S and his-FUSLC were used in control studies. Each protein was fluorescently labeled at amine groups using an N-hydroxysuccinimide (NHS)-reactive dye, Atto 488 for visualization, as described under Materials and Methods. The protein to Atto 488 ratio was less than 1:1. We observed the impact of each of the three proteins on membrane shape by incubating the proteins with giant unilamellar vesicles (GUVs) containing DOGS-Ni-NTA lipids. GUVs consisted of 83 mol% POPC, 15 mol% DGS-NTA-Ni, 2 mol% DP-EG10 biotin for coverslip tethering and 0.1 mol% Texas Red-DHPE for visualization.

**Figure 1.**
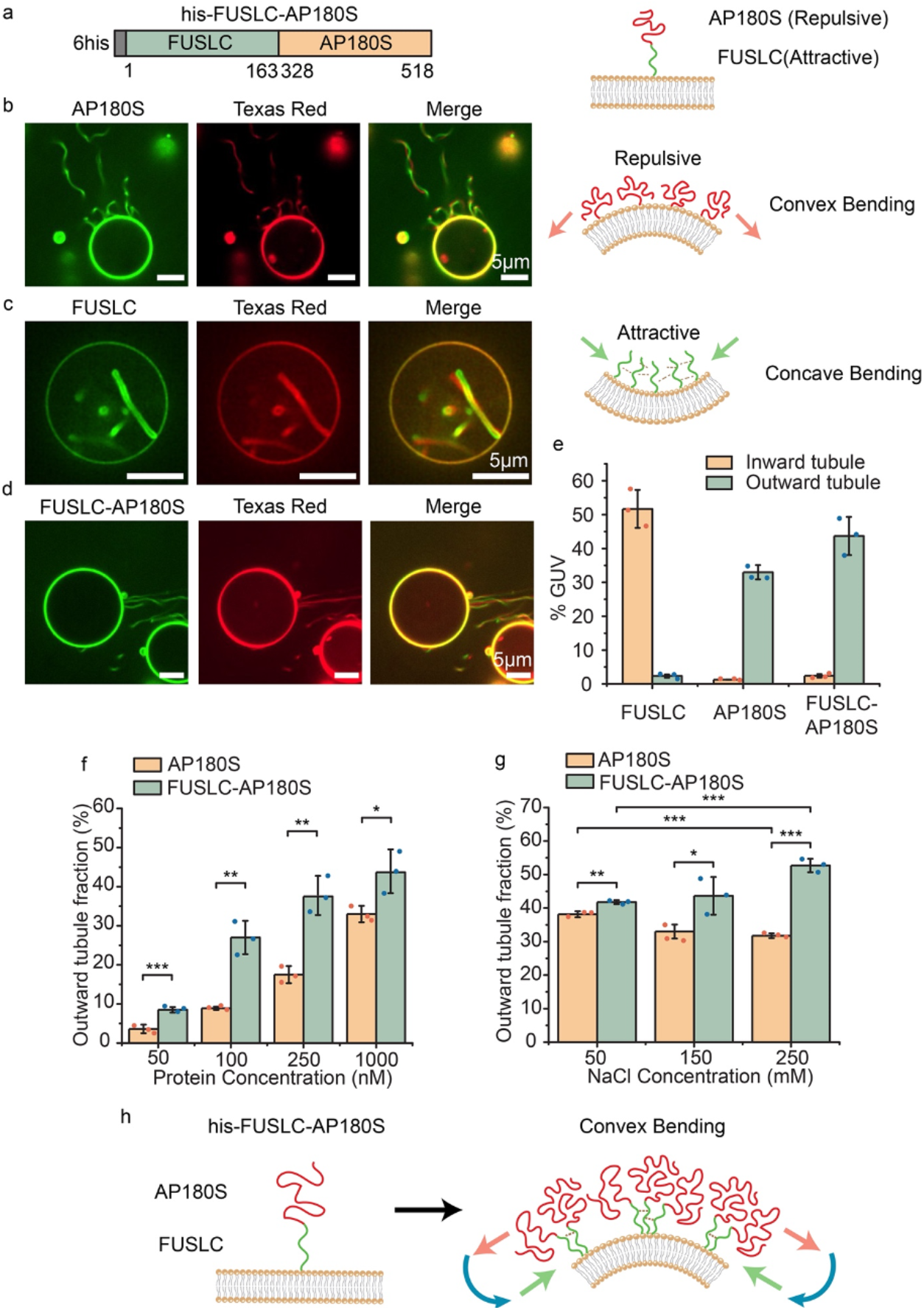
Attractive domains cluster repulsive domains at membrane surfaces, amplifying convex membrane bending. (a) Schematic of his-FUSLC-AP180S (left) and the expected orientation of the two domains relative to the membrane surface, when the histidine tag binds Ni-NTA lipids (right). (b) Representative images of protein-coated tubules emanating from GUV surfaces (protein and lipid channels) when incubated with 1μM his-AP180S (left). Cartoon of convex bending by repulsive intrinsically disordered proteins (IDPs) (right). (c) Representative super-resolution images of protein-lined tubules emanating from GUV surface when incubated with 1 μM his-FUSLC (left) and the corresponding cartoon of concave membrane bending by IDPs that attract one another (right). (d) Representative images of outward tubule formation when 1 μM his-FUSLC-AP180S was applied to GUVs. All scale bars are 5 μm. (e) The fraction of GUVs displaying inward and outward tubules when incubated with 1 μM his-FUSLC, his-AP180S, or his-FUSLC-AP180S. (f, g) The fraction of GUVs displaying outward tubules as a function of protein concentration (f), and under different NaCl concentrations when incubated with 1 μM protein (g). Error bars represent the standard deviation of three independent trials (displayed by the dots). Significance was evaluated using an unpaired, two-tailed student’s t test. *: P < 0.05, **: P < 0.01, ***: P < 0.001. Significance comparison between inward and outward tubule fraction for FUSLC, AP180S, and FUSLC-AP180S in panel (e) all have p value smaller than 0.001. GUV membrane composition was 83 mol% POPC, 15 mol% DGS-NTA-Ni, 2 mol% DP-EG10 biotin, and 0.1 mol% Texas Red-DHPE. All experiments were conducted in 25 mM HEPES, 150 mM NaCl buffer, pH 7.4 unless the NaCl concentration was specifically adjusted as shown in individual panels. (f) Schematic of attractive interactions among FUSLC domains amplifying crowding and repulsion among AP180S domains, leading to convex membrane bending.

When GUVs were exposed to 1 μM of his-AP180S, we observed protein recruitment to GUV surfaces within minutes, followed by emergence and extension of lipid tubules directed outward from the surfaces of GUVs (Figure 1b). Consistent with our previous reports, these tubules were diffraction limited in width and had lengths that often approached or exceeded the initial diameter of the GUVs(*7, 16, 17*). The tubules were visible in both the protein (Atto 488) and lipid (Texas Red) fluorescent channels. Because the protein was added to the outside of the GUVs and was excluded from the GUV lumens (Figure S1), we inferred that the protein must coat the outer surfaces of these convex tubules, as we have reported previously(*7*). Approximately 33% of GUVs exposed to his-AP180S displayed outwardly directed membrane tubules, while inwardly directed tubules were observed very rarely (Figure 1e).

In contrast, when GUVs were exposed to 1 μM of his-FUCLC, the protein was recruited to GUV surfaces within minutes, followed by emergence of inwardly directed membrane tubules. As we have reported previously(*12*), these tubules often displayed undulating morphologies, with diameters ranging from a few hundred nanometers to micrometers, such that the lumens of the tubules could often be resolved by fluorescence microscopy with deconvolution, Figure 1c. As with tubules formed upon addition of his-AP180S, tubules formed upon addition of his-FUSLC colocalized in the membrane and protein fluorescent channels. Owing to exclusion of protein from the GUV lumen, and the inward direction of the tubules, we inferred that the his-FUSLC protein lined these concave tubules(*12*). Approximately 52% of GUVs exposed to his-FUSLC displayed inwardly directed membrane tubules, while outwardly directed tubules were observed very rarely (Figure 1e). Notably, outwardly directed tubules were generally narrower than inwardly directed tubules, likely because the attractive interactions between proteins that drive inward tubules tend to simultaneously increase the membrane rigidity(*12*), making it more difficult to curve the membrane.

When GUVs were exposed to 1 μM of the chimera, his-FUSLC-AP180S, it bound rapidly to the membrane surface, similar to the control proteins. Shortly after binding to the membrane surface, outwardly directed, protein-coated tubules were observed on GUV surfaces (Figure 1d), similar in morphology to those created by binding of his-AP180S (Figure 1b). Quantification of the frequency with which outwardly-directed tubules were observed revealed that the chimera, his-FUSLC-AP180S, was more likely to generate tubules when applied at a given solution concentration, in comparison to his-AP180S (Figure 1f). Increasing the concentration of sodium chloride (NaCl) in the buffer slightly decreased the formation of outwardly directed membrane tubules by his-AP180S, presumably by screening electrostatic repulsion, as described previously(*14*). In contrast, the same increase in NaCl concentration somewhat increased formation of outwardly directed tubules by the chimera, his-FUSLC-AP180S, Figure 1g. This trend suggests that clustering of FUSLC domains, which increases with increasing NaCl concentration(*15*), promotes outward tubule formation by the chimera. Notably, changes in pH might also be capable of shifting the balance between attractive and repulsive interactions among the chimeras. However, the substantial shifts in pH that would be required to change the net charge of the chimeras are likely to also change the mechanical properties of the membranes(*18*), making the results difficult to interpret. Collectively, these results suggest that the presence of the FUSLC domain enhanced formation of outward tubules by the AP180S domain, perhaps by forming local clusters of the protein, which would be expected to enhance membrane binding, helping to generate local steric pressure (Figure 1h), which is then relaxed by membrane bending. In particular, when FUSLC domains bind to one another, their associated AP180S domains are brought into close contact with one another, resulting in a local increase in the density of AP180 domains on the membrane surface. Because steric pressure is expected to increase non-linearly with increasing density of AP180 domains(*7*), the close contact created by association between the FUSLC domains is likely responsible for the increased capacity of his-FUSLC-AP180S to generate protein-coated membrane tubules in comparison to his-AP180S(*8*).

These results suggest that the preference of the AP180S domain for convex curvature dominate over the preference of FUSLC for concave curvature. This dominance could result simply from the magnitude of the repulsive interactions generated by AP180S exceeding the magnitude of attractive interactions generated by FUSLC. Alternatively, the dominance of AP180S could arise from its position further from the neutral surface of the curved membrane, such that repulsive interactions among AP180S domains generate a larger bending moment in comparison to attractive interactions among FUSLC domains, as depicted in Figure 1h. Based on these results alone, it is unclear to what extent the order of the protein domains relative to the membrane surface plays a role in curvature generation.

### Reversing the order of the domains relative to the membrane surface reverses the direction of membrane curvature

Based on our findings in Figure 1, we next asked what might happen if we reversed the order of the domains, creating a chimera that combined repulsive interactions at the membrane surface with attractive interactions farther away. To answer this question, we created the chimera his-AP180S-FUSLC, Figure 2a. When this chimera, labeled with Atto488 dye, bound to the surfaces of GUVs, we observed phase separation of the protein on the membrane surface, resulting in rounded, spherical cap-like structures on the membrane surface, which were enriched in the chimeric protein, Figure 2b. These structures were similar to those that we observed previously with his-FUSLC(*12*), suggesting that the his-AP180S-FUSLC chimera underwent coacervation on membrane surfaces. Within minutes after protein binding, many of the protein-rich regions spontaneously curved inward, creating protein-lined membrane tubules with similar morphologies to those observed upon exposure of vesicles to his-FUSLC, Figure 2c. The fraction of GUVs displaying protein phase separation and protein-lined tubules increased significantly as salt concentration increased (Figure 2d), suggesting that the attractive interaction became stronger at higher salt concentration, which is consistent with the behavior of FUSLC alone(*12, 15*). Similarly, the diameters of the tubules formed by the his-AP180S-FUSLC chimera were typically resolvable using deconvoluted confocal fluorescence microscopy. Interestingly, the distribution of tubule diameter shifted toward larger values as the concentration of NaCl increased, Figure 2e. This shift may be due to increased attraction among FUSLC domains with increasing salt concentration(*15*), which may increase the rigidity of the protein layer on the membrane surface, making it more difficult for the membrane to take on high curvature(*12*). Notably, the fraction of vesicles displaying phase separation, the fraction of vesicles displaying inward tubules, and the average diameter of the tubules were all significantly lower than the corresponding values for vesicles exposed to FUSLC alone (Figure S2)(*12*). These results suggest that AP180S may weaken the attractive interactions among FUSLC domains, such that phase separation is less effective, leaving the membrane more flexible, such that tubules of higher curvature can be formed.

**Figure 2.**
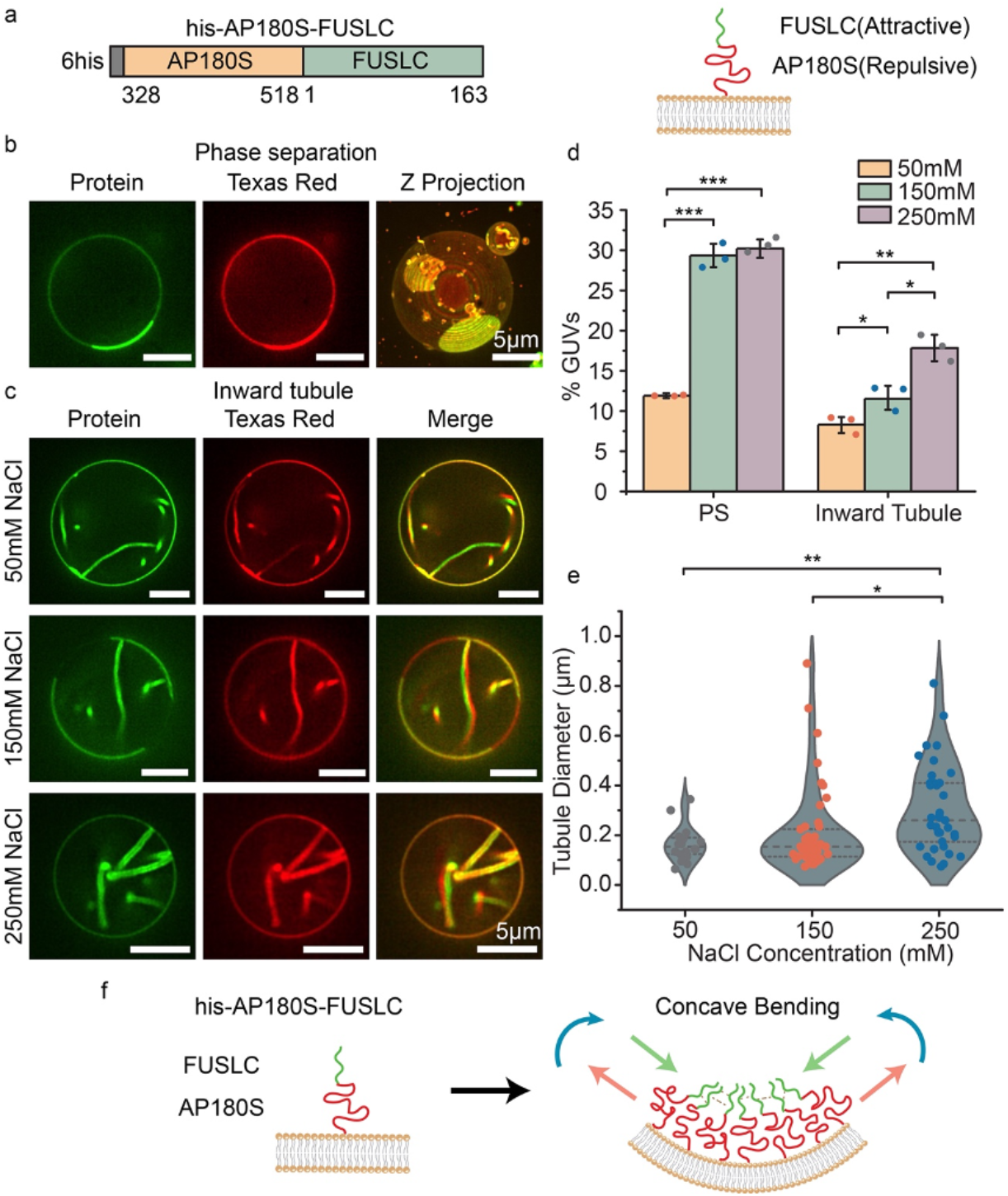
Using a repulsive domain to link an attractive domain to the membrane surface generates protein-lined, concave membrane tubules. (a) Schematic of the recombinant chimera his-AP180S-FUSLC (left panel) and the orientation of his-AP180S-FUSLC on the membrane surface when it binds to DGS-NTA-Ni lipids (right panel). (b) Representative super-resolution images (protein and lipid channel) of protein liquid-liquid phase separation when GUVs were exposed to 1 μM his-AP180S-FUSLC in 25 mM HEPES, 150 mM NaCl buffer, pH 7.4. (c) Representative protein and lipid channel confocal images of tubules emanating inward from GUV surfaces when incubated with 1 μM his-AP180S-FUSLC in 25 mM HEPES, pH 7.4 buffer containing 50 mM, 150 mM, and 250 mM NaCl, respectively. Scale bar is 5 μm. (d) Fraction of GUVs showing protein phase separation (PS) and inward tubules in the presence of different NaCl concentrations. (e) Violin plot displaying inward tubule diameter distribution in the presence of different NaCl concentrations. Error bars represent the standard deviation from three independent trials (shown by the dots). In total, n >100 GUVs were imaged for each NaCl concentration. Statistical significance was tested using an unpaired, two-tailed student’s t test. *: P < 0.05, **: P < 0.01, ***: P < 0.001. GUV composition was 83 mol% POPC, 15 mol% DGS-NTA-Ni, 2 mol% DP-EG10 biotin, and 0.1 mol% Texas Red-DHPE. (f) Schematic of concave membrane bending when attractive domains are further from the membrane surface, relative to repulsive domains.

Collectively, these results suggest that attractive interactions among FUSLC domains at a distance from the membrane surface drive the membrane to bend inward, generating protein-lined tubules. The observation that the two chimeras, his-FUSLC-AP180S and his-AP180S-FUSLC generate tubules of opposite curvature suggests that the two chimers create stresses with opposite signs on the membrane surface. In particular, the observation of convex, outwardly directed tubules, suggests that his-FUSLC-AP180S stretches the outer membrane leaflet while compressing the inward leaflet, generating a bending moment orientated as shown in Figure 1h. This orientation is consistent with attractive forces closer to the membrane surface and repulsive forces farther away, as depicted in the figure. In contrast, the observation of concave, inwardly directed tubules, suggests that his-AP180S-FUSLC compresses the outer membrane leaflet while stretching the inward leaflet, generating a bending moment that is oriented as shown in Figure 2f, opposite to that in Figure 1h. This orientation is consistent with repulsive forces adjacent to the membrane surface and attractive forces farther away, as indicated in the figure. Both results suggest that AP180S and FUSLC form somewhat separate layers on the membrane surface, which is consistent with the relative exclusion of AP180S from droplets consisting of FUSLC (Figure S3). Collectively, these results demonstrate that the orientation of disordered protein domains relative to the membrane surface can be used to control the magnitude and direction of the bending moment they exert, ultimately providing control over membrane shape.

### Combining chimeras with opposite impacts on membrane curvature provides control over membrane shape

If his-FUSLC-AP180S and his-AP180S-FUSLC apply opposite bending stresses on the membrane surface, then it should be possible to control the direction of membrane bending by exposing vesicles to varying ratios of the two chimeras (Figure 3a). To evaluate this prediction, we exposed giant vesicles (unlabeled) to his-AP180S-FUSLC, labeled with Atto488 (green), and his-FUSLC-AP180S, labeled with Atto594 (red) (Figure 3a, b). his-FUSLC-AP180S and his-AP180S-FUSLC were combined in ratios ranging from 0.1: 1 to 1:1. At the lowest ratios, where his-AP180S-FUSLC dominated, vesicles with inwardly-directed tubules were most common (Figure 3c top panel). In contrast, for the highest ratios, where the two chimeras had equal concentrations, outwardly directed tubules dominated (Figure 3c bottom panel). For the intermediate ratio of 0.25:1, inward and outward tubules each existed in about 5% of vesicles separately, with the remaining 90% of membranes lacking tubules (Figure 3c middle panel and Figure 3d). In all cases, it was very rare to observe vesicles with both inward and outward tubules present simultaneously. Instead, nearly all vesicles exhibited tubules of a single orientation, (Figure 3d). These results suggest that the bending moments generated by the two chimeras can be balanced out when they are combined, stabilizing a relatively flat membrane morphology. This balance is likely enabled by the ability of the two chimeras to mix with one another when they bind to membrane surfaces. At all ratios, we observed colocalization between the fluorescence signals associated with the two proteins, suggesting that they were not segregated on the membrane surface. This mixing behavior is expected, as both chimeras contain the FUSLC domain. Nonetheless, inward tubules appeared somewhat enriched in the chimera that prefers concave curvature, his-AP180S-FUSLC (Figure 3c, 0.1:1, S4), while outward tubules appeared somewhat enriched in the chimera that prefers convex curvature, his-FUSLC-AP180S (Figure 3c, 0.5:1). Taken together, these results illustrate that protein domains with opposite curvature preferences can work together to maintain flat membrane surfaces.

**Figure 3.**
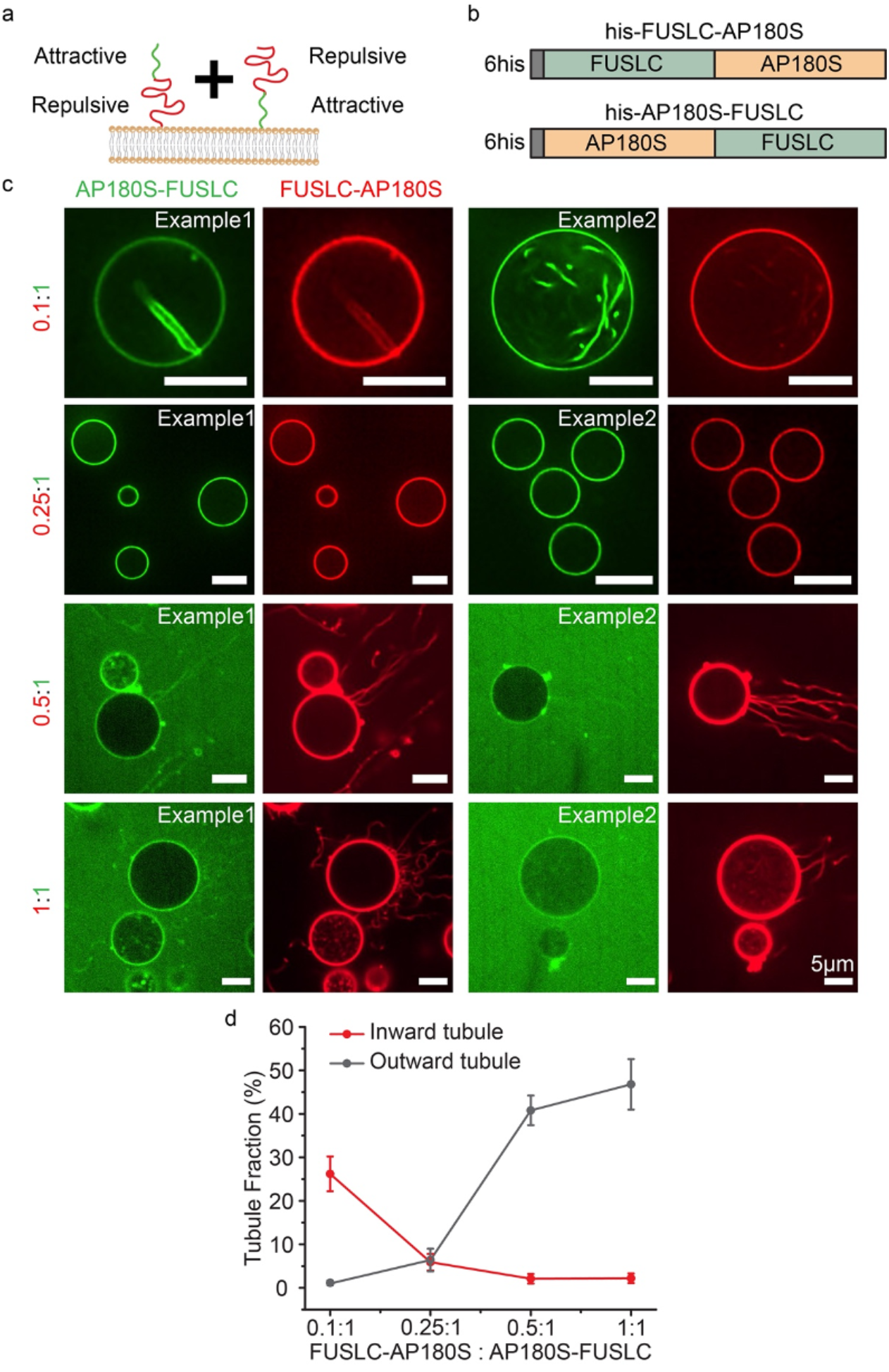
Combining chimeras with opposite impacts on membrane curvature provides control over membrane shape. (a) Cartoon of adding his-FUSLC-AP180S and his-AP180S-FUSLC simultaneously to the membrane and the relative position of the attractive and repulsive domains relative to the membrane surface. (b) Schematic of the two recombinant chimera his-FUSLC-AP180S and his-AP180S-FUSLC. (c) Representative confocal images of GUVs incubated with his-FUSLC-AP180S and his-AP180S-FUSLC, mixed at different ratios (from 0.1:1 to 1:1). His-AP180S-FUSLC was maintained at 1 μM in all conditions. Experiments were done in 25 mM HEPES, 150 mM NaCl buffer, pH 7.4. GUV composition was 83 mol% POPC, 15 mol% DGS-NTA-Ni, 2 mol% DP-EG10 biotin, and 0.1 mol% Texas Red-DHPE. All scale bars are 5 μm. (d) The frequency of GUVs displaying outward tubules and inward tubules as a function of his-FUSLC-AP180S to his-AP180S-FUSLC ratio. Error bars represent the standard deviation from three independent trials. In total n >200 GUVs were imaged for each ratio.

### Ionic strength can shift the balance between attractive and repulsive interactions, reversing the direction of membrane curvature

If membrane curvature results from the balance between attractive and repulsive interactions among disordered domains, then it should be possible to modulate curvature by perturbing this balance. We tested this principle by using changes in ionic strength to vary the relative magnitude of attractive and repulsive interactions. Specifically, to achieve a greater dynamic range in the magnitude of the repulsive interactions, we created a new chimera, which linked FUSLC to the full C-terminal domain of AP180 (residues 328-898), which has a net negative charge of -32(*14*). We will refer to this domain as the “long version” of AP180, or AP180L, yielding the chimera, his-AP180L-FUSLC, Figure 4a, b. By incorporating a larger portion of AP180, with a larger hydrodynamic radius(*14*), this chimera should generate a larger steric pressure, which may be capable of overcoming the attractive interactions among the FUSLC domains, depending on the ionic strength. It is worth noting that the effect of his-FUSLC-AP180L was not tested, since, on the basis of our findings with his-FUSLC-AP180S, we would simply expect more outward tubules if we were to make the repulsive domain larger.

**Figure 4.**
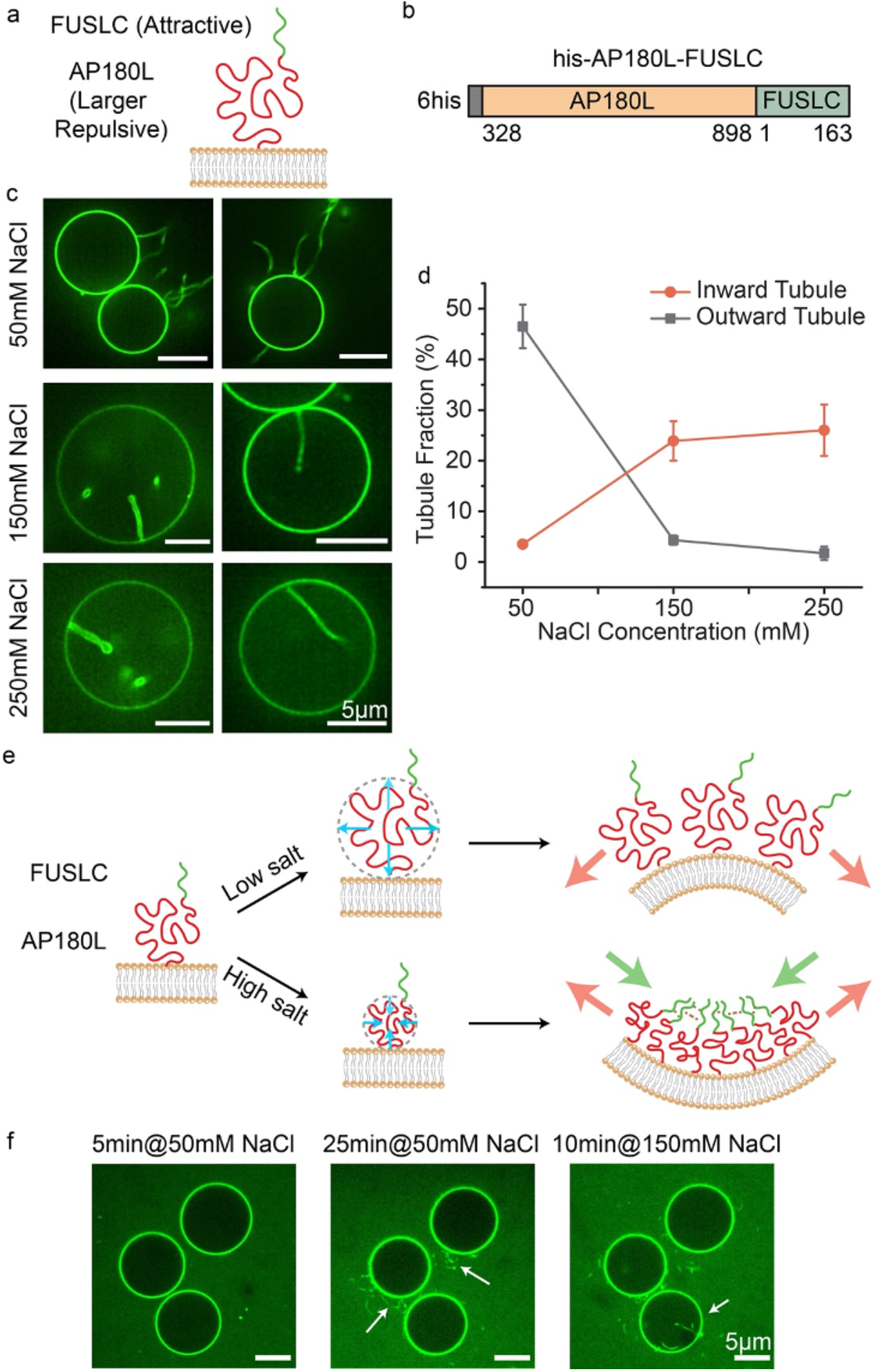
Ionic strength can shift the balance between attractive and repulsive interactions, reversing the direction of membrane curvature. (a,b) Schematic of the orientation of his-AP180L-FUSLC when binding to the membrane (a), and the diagram of the domains (b). (c) Representative deconvoluted images of GUVs when incubated with 1 μM his-AP180L-FUSLC in 25 mM HEPES, pH 7.4 buffer containing 50 mM, 150 mM, and 250 mM NaCl, respectively. Scale bar is 5 μm. (d) The frequency of GUVs displaying outward tubules and inward tubules as a function of NaCl concentration. Error bars represent the standard deviation of three trials. n > 90 GUVs were imaged in each trial. GUV composition was 83 mol% POPC, 15 mol% DGS-NTA-Ni, 2 mol% DP-EG10 biotin, and 0.1 mol% Texas Red-DHPE. (e) Schematic depicting dependence of membrane curvature on ionic strength. (f) *In-situ* observation of outward and inward tubule formation as salt concentration increases.

Giant vesicles were exposed to 1 μM of his-AP180L-FUSLC at a range of NaCl concentrations: 50 mM, 150 mM, and 250 mM, Figure 4c, d. At 50 mM NaCl, exposure to the chimera drove formation of protein-coated, outwardly directed membrane tubules. This result suggests that electrostatic repulsion among the AP180L domains, which is maximized at low NaCl concentration, dominated over attractive interactions between FUSLC domains, setting up an outwardly directed bending moment, as shown in Figure 4e, top. In contrast, at the higher salt concentrations, 150 mM and 250 mM, exposure to the chimera drove formation of protein-lined, inwardly directed tubules, suggesting that repulsive interactions among the AP180L domains, which are reduced at higher salt concentrations, were overcome by attractive interactions among FUSLC domains, setting up an inwardly directed bending moment, as shown in Figure 4e, bottom. Further, we were able to observe the transition from outward to inward tubules *in-situ* as we gradually changed the NaCl concentration from 50 mM to 150 mM (Figure 4f). Collectively these results demonstrate that it is possible to control the direction of membrane bending by changing environmental conditions, such as ionic strength, which alter the balance between attractive and repulsive interactions among disordered protein domains. Depending on the domains used, other environmental variables such as pH, temperature, and the presence of multivalent ligands could also be used to tune interaction strength.

### Reversing the response of the attractive domain to ionic strength reverses the direction of membrane tubules

Having demonstrated the ability to control the direction of membrane curvature by varying ionic strength, we next sought to test the generality of the principle by altering the response of the attractive domain to NaCl concentration. Specifically, we replaced FUSLC with the RGG domain (1-168) of the Laf-1 protein(*19, 20*), another disordered domain that is known to undergo liquid-liquid phase separation. RGG is rich in positively charged arginine residues and negatively charged aspartic acid residues, such that electrostatic attraction plays a major role in the coacervation of RGG domains(*10*). For this reason, attraction between RGG domains is expected to decrease with increasing NaCl concentration, opposite to the response of FUSLC.

To test the impact of replacing FUSLC with RGG, we constructed a chimera between AP180L and RGG, his-AP180L-RGG (Figure 5a). We exposed giant unilamellar vesicles to these chimeras at a concentration of 1 μM, while gradually increasing the NaCl concentration. At the lowest concentrations of NaCl, inwardly directed tubules were much more probable in comparison to outwardly directed tubules, which were rarely observed (Figure 5b). This result suggests that electrostatic interactions among RGG domains, which are strongest at low salt concentration, dominated over repulsive interactions among AP180L domains, leading to an inwardly directed bending moment, similar to what was observed for the his-AP180L-FUSLC chimera at high salt concentration.

**Figure 5.**
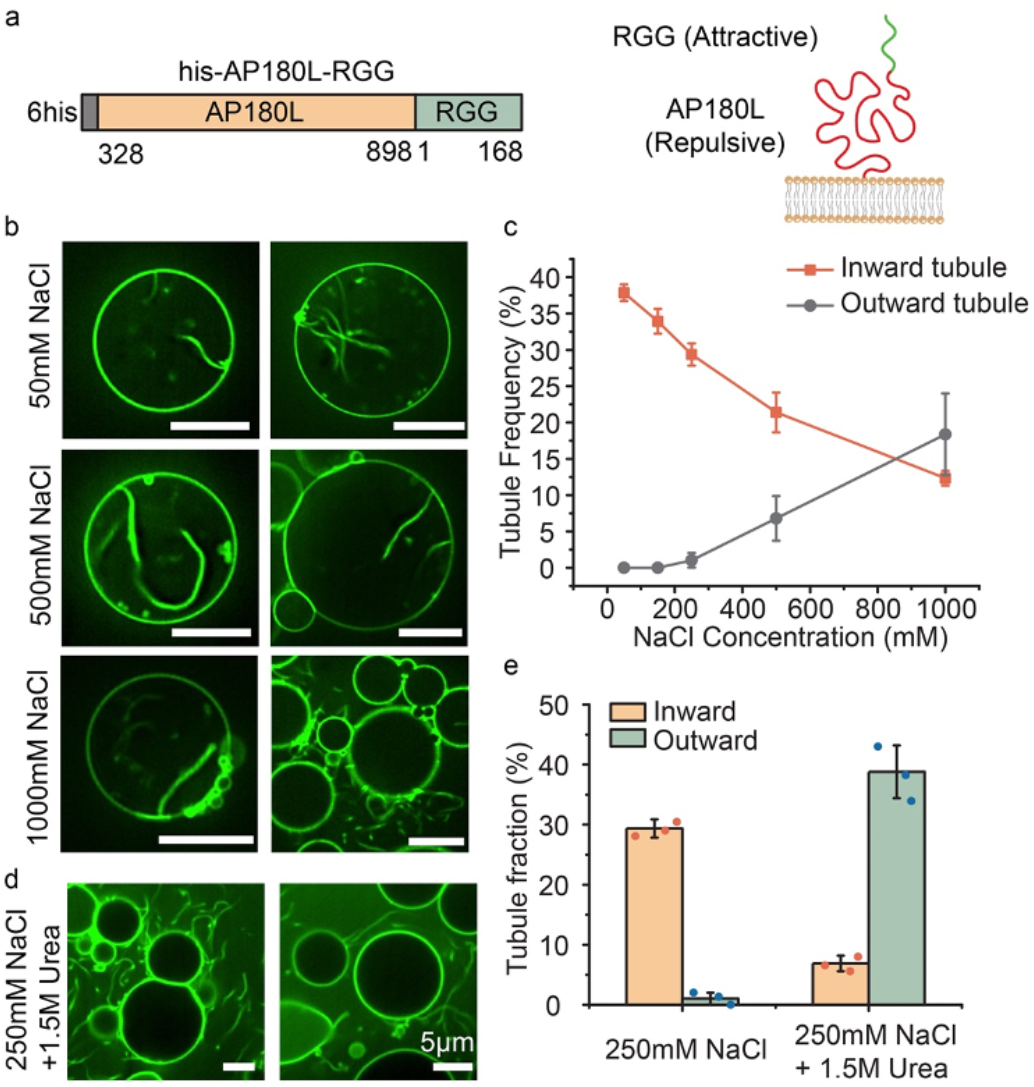
Reversing the response of the attractive domain to ionic strength reverses the direction of membrane tubules. (a) Schematic of his-AP180L-RGG domains (left panel) and their orientation relative to the membrane (right panel). (b) Representative confocal images of tubules emanating from GUVs incubated with 1 μM his-AP180L-RGG in 25 mM HEPES, pH 7.4 buffers containing 50 mM, 500 mM, or 1000 mM NaCl, respectively. (c) The fraction of GUVs displaying inward and outward tubules as a function of NaCl concentration. Data represent mean ± standard deviation, n = 3 independent experiments. (d) Representative confocal images of tubules emanating from GUVs incubated with 1 μM his-AP180L-RGG in 25 mM HEPES, 250 mM NaCl, pH 7.4 buffer with 1.5 M urea. (e) The fraction of GUVs exhibiting inward and outward tubules after exposure to 1 μM his-AP180L-RGG in 25 mM HEPES, 250 mM NaCl pH 7.4 buffer and 25 mM HEPES, 250 mM NaCl pH 7.4 buffer with 1.5 M urea. Error bars represent the standard deviation of three independent trials (indicated by the dots). Statistical significance between inward and outward tubule frequency was tested by an unpaired, two-tailed student’s t test. In panel (e) both have p values were smaller than 0.001. n > 100 GUVs were imaged in each trial. GUV composition was 83 mol% POPC, 15 mol% DGS-NTA-Ni, 2 mol% DP-EG10 biotin, and 0.1 mol% Texas Red-DHPE. (Scale bars in b and d are 5 μm.)

As the NaCl concentration increased, the frequency of inwardly directed tubules fell as the frequency of outwardly directed tubules increased. Only at 1 M NaCl did outwardly directed tubules become more frequent than inwardly directed tubules (Figure 5c). As predicted, this trend is opposite to that observed for his-AP180L-FUSLC, where outwardly directed tubules dominated at low salt concentrations. Interestingly, when we added urea (at 250 mM NaCl) to attenuate all protein-protein interactions, we mainly observed outward tubules, as would be expected when non-specific, steric interactions are dominant. This result is the opposite of what we observed at the same NaCl concentration in the absence of urea (Figure 5d, e), further confirming that specific, attractive interactions among RGG domains provided the driving force for inward bending of the membrane. Collectively these results demonstrate that by reversing the response of the attractive domain to changes in ionic strength, it is possible to reverse the sign of the bending moment that the protein layer applies to the membrane, resulting in a reversal of the direction of membrane curvature.

### A simple mechanical model reproduces the ability of disordered protein chimeras to drive inward and outward curvature

In Figure 6a, we summarize our experimental results in the form of an empirical phase diagram, in which the curvature driving abilities of the chimeras are represented as a function of salt concentration and relative separation between the attractive, LLPS-inducing domain (FUSLC or RGG), and the membrane surface. Here we see that the domain that is farthest from the membrane surface generally determines whether the net interaction between chimeras is attractive or repulsive. For example, the his-FUSLC-AP180S chimera, which places the FUSLC domain in closest proximity to the membrane surface, resulted in outward membrane tubules at all salt concentrations, suggesting that repulsion among AP180S domains dominated. In contrast, the his-AP180S-FUSLC chimera, which places the FUSLC domain farther from the membrane, resulted only in inward tubules, suggesting that attraction among the FUSLC domains dominated. Mixtures of these two chimeras effectively neutralized membrane curvature (blue circle). The his-AP180L-FUSLC chimera moves the FUSLC domain even farther from the membrane surface, but also increases the potential for steric and electrostatic repulsion, by increasing the size of the AP180-derived domain. As a result, the impact of this chimera on membrane curvature was conditional, with electrostatic repulsion among AP180L domains dominating at low salt concentration, resulting in outward tubules, and attraction among FUSLC domains dominating at higher salt concentrations, resulting in inward tubules. As described in the previous section, the his-AP180L-RGG chimera (not shown on the phase diagram) reversed this trend, owing to the opposite response of RGG-RGG interactions to changes in salt concentrations, when compared to FUSLC-FUSLC interactions(*10*).

**Figure 6.**
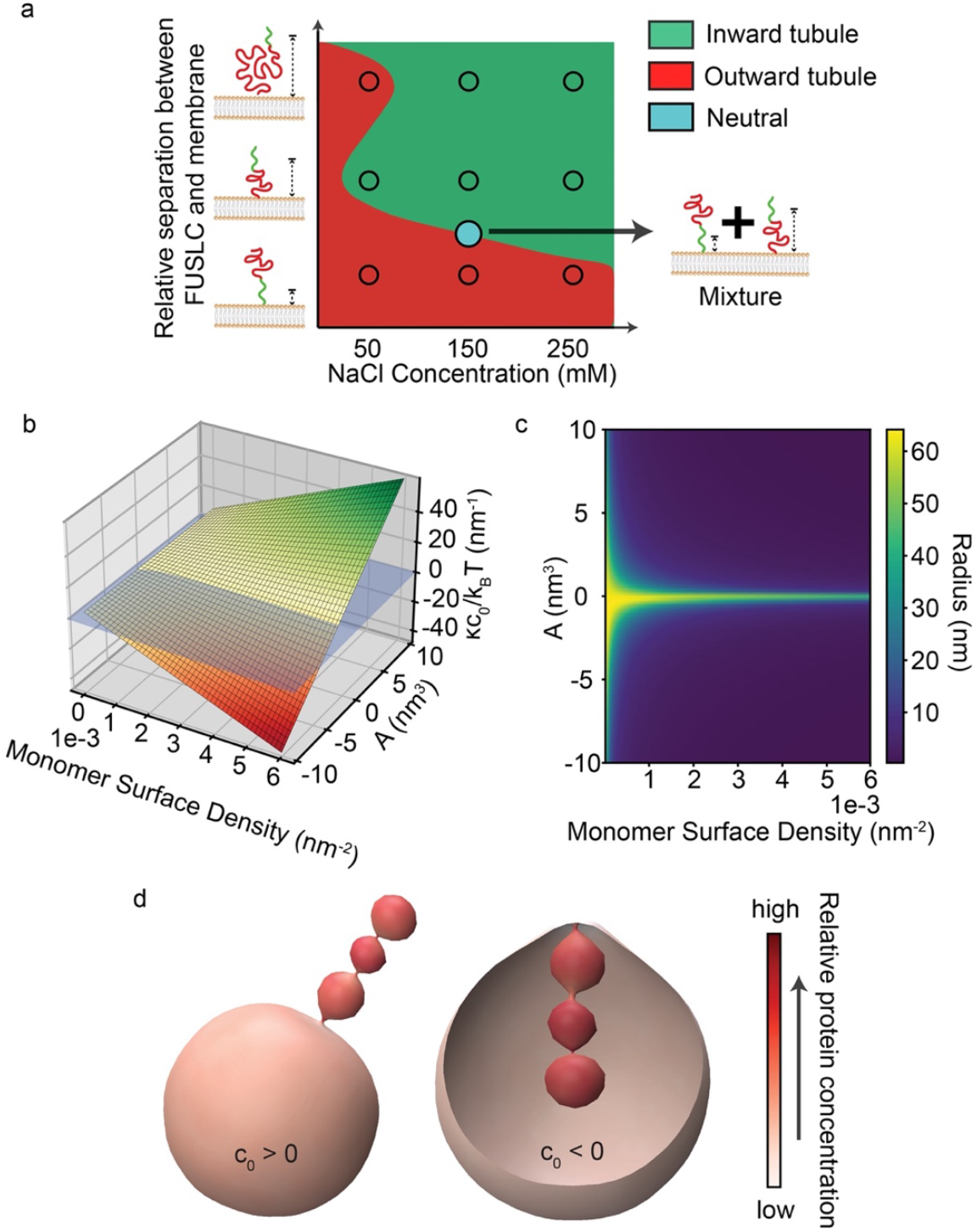
A simple mechanical model reproduces the ability of disordered protein chimeras to drive inward and outward curvature. (a) Empirical phase diagram displaying the membrane curvature preferences of disordered protein chimeras as a function of salt concentration and distance of the attractive domain (FUSLC) from the membrane surface. (b) Modeling the membrane attached chimeras as a polymer brush demonstrates that the bending moment induced by the proteins (vertical axis) increases with increasing concentration of monomers at the membrane surface and second virial coefficient, *A*. (c) Predicted tube radius as a function of monomer surface density and *A*. (d) Simulation of inward and outward tubulation using Mem3DG. The left and right renderings simulate the impact of proteins with positive and negative spontaneous curvatures, respectively. Color indicates local protein concentration, demonstrating how proteins accumulate and reinforce the curvature of the tubule.

Taken together, these data suggest that when the net interaction between the domains that make up a chimera is attractive, the chimera will drive inward membrane bending. In contrast, if the net interaction is repulsive, the chimera will drive outward bending. To illustrate this concept, we developed a simple mechanical model, which treats the layer of membrane-bound protein chimeras as a layer of polymers chains grafted onto a flexible surface (Figure 6b-d, S5), see Supplementary Information. Briefly, following recently published theoretical work on membrane-bound polymer brushes(*9, 21-24*) we wrote an expression for the free energy of the protein-membrane composite system (Equations S1, S2). The free energy of the brush-like IDP layer includes contributions from chain stretching, attractive or repulsive interactions between chains, and the loss of entropy upon ion partitioning into the brush. The membrane energy is the conventional Helfrich-Canham-Evans Hamiltonian(*25-27*). Minimizing the free energy with respect to membrane curvature, Figure 6b plots the equilibrium membrane bending moment (vertical axis), which is proportional to spontaneous curvature, as a function of the density of protein monomers at the membrane surface, and the second virial coefficient associated with chain-chain interactions, *A* (horizontal axes). *A* is positive when the chains have net repulsive interactions and negative when their net interactions are attractive. In agreement with our experimental results, this simple model predicts that membrane curvature will increase with the monomer density and will be positive (outward curvature) when the chains repel one another and negative (inward curvature) when they attract one another. With the range of spontaneous curvatures obtained in Figure 6b, we next minimized the free energy of a cylindrical tube and found the corresponding tube radii as a function of monomer concentration and *A*, Figure 6c.

To demonstrate inward and outward tube formation, we used Mem3DG(*28*) to build a model of the membrane surface in three dimensions. Mem3DG is a framework and simulation engine that enables us to solve the governing equations of membrane bending using principles of discrete differential geometry. For this demonstration, we started from a spherical vesicle and applied a weak Gaussian point force, decaying in time, to induce initial tube formation. Proteins which impart spontaneous curvature matching the direction of the tube bind and support the extrusion of the tubule. The resulting configurations, shown in Figure 6d, show that proteins that impart a positive spontaneous curvature stabilize outward tubules (left), while proteins that impart a negative spontaneous curvature stabilize inward tubules (right), suggesting that this simple physical model captures the main features of our experimental system. In both cases, the tubules have a pearled morphology similar to those observed in experiments, (Figure 2c, 4c).

## Discussion

In this study, we have shown that the balance between attractive and repulsive interactions determines the extent and direction of membrane curvature by intrinsically disordered proteins. Specifically, while disordered domains that repel one another generate steric pressure that drives outward, convex membrane bending^7^, attractive interactions among disordered domains do the opposite, generating compressive stress that results in inward, concave membrane bending(*12*). By generating chimeras that combined these two types of interactions within the same protein, we have illustrated a set of design principles that can be used to control membrane shape in response to external stimuli.

The first principle is that the orientation of a chimera with respect to the membrane surface can be used to control the direction of membrane bending. Specifically, the domain farthest from the membrane surface tends to dominate the membrane curvature, likely because the farther domain has the advantage of a larger moment arm, and can therefore apply a larger bending stress. In this way, putting the repulsive domain farther from the membrane surface led to convex, outward bending, while placing the attractive domain farther from the membrane surface led to concave, inward bending (Figure 1d, 2c).

The second principle we identified is that chimeras with opposite curvature preferences can be used to counteract one another, leading to a relatively flat, stable membrane even in the presence of these proteins. In particular, we mixed the convex curvature preferring chimera (his-FUSLC-AP180S) with the concave preferring chimera (his-AP180S-FUSLC) at a range of ratios, and identified an intermediate regime in which the membrane preferred to remain flat, whereas tubules, of either convex or concave curvature, dominated at other ratios (Figure 3c). Flat, protein-decorated, membranes are found throughout the cell, from regions of the plasma membrane to the sheets of the endoplasmic reticulum and the cisternae of the Golgi apparatus(*1, 2, 29, 30*), suggesting the importance of stabilizing flat, as well as curved, membrane shapes in the presence of membrane-bound proteins.

The final design principle we identified is that membrane curvature can be tuned and even reversed when the balance between attractive and repulsive protein interactions is altered by environmental changes. Specifically, we used a decrease in ionic strength to simultaneously strengthen electrostatic repulsion between AP180L domains, while weakening attractive interactions among FUSLC domains. These collective effects ultimately reversed the direction of membrane bending. Specifically, the chimeric protein his-AP180L-FUSLC formed concave tubules at high ionic strength and convex tubules at low ionic strength (Figure 4c). This result illustrates that the dominance of the domain farthest from the membrane surface – identified in the first principle – can ultimately be overcome if there is a sufficient imbalance between attractive and repulsive interactions. Sensitivity of membrane-protein interactions to changes in the local environment is likely an important factor in cellular and organellar membranes where curvatures of both directions are observed, such as the endoplasmic reticulum and the inner membrane of mitochondria.

Collectively, these principles illustrate that the diverse and dynamic curvatures found in cellular membranes can be achieved by disordered proteins, entirely in the absence of structured domains. As 30-50% of all proteins are now thought to contain significant regions of intrinsic disorder(*31, 32*), these observations have the potential to substantially expand our understanding of the proteome responsible for membrane curvature. More broadly, many proteins that are known to play a role in defining membrane shape contain both structured and intrinsically disordered domains(*33*). On the basis of our current and previous findings(*8*), it appears increasingly likely that structured and disordered curvature drivers collaborate to define the shape of cellular membranes. As one example, recent work has illustrated that BAR domains, structured curvature-inducing scaffolds, are often found within proteins that also contain substantial disordered domains, such that steric pressure among the disordered domains amplifies the inherent curvature preference of the BAR domains, resulting in convex membrane curvature(*8*). Another example is the influenza matrix protein M1, which provides the major driving force in virus budding(*34*). A recent *in vitro* study showed that the structured N terminal domain of M1 binds to the membrane, but requires the disordered C terminal domain to achieve polymerization, ultimately driving concave membrane invagination(*35*). Coordination between structured and disordered domains may also play a role in maintaining the curvature of the nuclear pore complex, which is lined by nucleoporins that contain disordered domains rich in phenylalanine-glycine (FG) repeats(*36-38*). Recent work suggests that the FG-rich domains form a flexible network that has the properties of a protein condensate(*39*). Based on our findings, interactions between these domains could help to stabilize the complex architecture of the nuclear pore, which contains both convex and concave curvatures. Inspired by these examples and the growing recognition of the role that disordered proteins play in curving membranes, the design rules identified in the present study have broad implications for our understanding of the diverse mechanisms by which protein networks shape biological membranes.

## Author Contributions

F.Y, J.R.H, E.M.L, P.R, and J.C.S designed the research. F.Y and A.S performed the cloning for all constructs. L.W and F.Y purified all the protein chimeras used in this research. F.Y and J.R.H performed the imaging. F.Y, J.R.H, P.R and J.C.S contributed to data analysis. C.T.L and P.R. performed the modeling and simulations. F.Y, L.W, E.M.L, C.T.L., P.R and J.C.S wrote the paper.

The authors declare no competing interest.

## Data Availability

All data needed to evaluate the conclusions in the paper are present in the paper and/or the Supplementary Materials.

## Materials Availability

Plasmids for bacterial expression of the chimeric proteins used in this work are available from the corresponding author upon request. Detailed protocols for purification of the chimeras are include in the materials and methods section.

## Acknowledgement

This research was supported by the NIH through grants R35GM139531 (J.C.S), R01GM132106 (P.R), the NSF DMS through grant 1934411 (P.R and J.C.S), ONR N00014-20-1-2469 to (P.R), the Welch Foundation through grant F-2047 (J.C.S.), Kavli Institute for Brain and Mind Postdoctoral Award (C.T.L) and the University of Texas at Austin, Graduate School Continuing Fellowship 2022-2023 (F.Y). The authors would like to thank Cuncheng Zhu and Arijit Mahapatra for helpful discussions.

## Materials and Methods

### Reagents

POPC (1-palmitoyl-2-oleoyl-glycero-3-phosphocholine) and DGS-NTA-Ni (1,2-dioleoyl-sn-glycero-3-[(N-(5-amino-1-carboxypentyl) iminodiacetic acid)-succinyl] (nickel salt)) were purchased from Avanti Polar Lipids, Inc. Texas Red-DHPE (Texas Red 1,2-dihexadecanoyl-sn-glycero-3-phosphoethanolamine triethylammonium salt), NeutrAvidin, TrisHCl (Tris hydrochloride), HEPES (4-(2-hydroxyethyl)-1-piperazineethanesulfonic acid), IPTG (isopropyl-β-D-thiogalactopyranoside), b-ME (b-mercaptoethanol), TCEP (tris(2-carboxyethyl)phosphine) and Triton X-100 were purchased from Thermo Fisher Scientific. Tryptone, Yeast Extract, NaCl, NaH_2_PO_4_, Na_2_HPO_4_, Urea, sodium tetraborate, EDTA (Ethylenediaminetetraacetic acid), CaCl_2_, glycerol, EDTA-free protease inhibitor tablets, imidazole, PMSF (phenylmethanesulfonylfluoride), PLL (poly-L-lysine), ATTO-594 NHS-ester, and ATTO-488 NHS-ester were purchased from Sigma-Aldrich. DP-EG10-biotin (dipalmitoyl-decaethylene glycol-biotin) was generously provided by Darryl Sasaki from Sandia National Laboratories, Livermore, CA(*40*). Amine reactive polyethylene glycol (mPEG-Succinimidyl Valerate MW 5000) and PEG-biotin (Biotin-PEG SVA, MW 5000) were purchased from Laysan Bio, Inc. All reagents were used without further purification.

### Plasmids

The DNA plasmids for AP180CTD (rat AP180, amino acids 328-898) in a pET32c vector was kindly provided by Ernst Ungewickell, Hannover Medical School, Germany. DNA coding for histidine-tagged AP180CTD (his-AP180CTD, denoted as his-AP180L) was cloned into a pGex4T2 vector as previously described(*7*) to incorporate a GST-tag at the N terminus of his-AP180CTD to stabilize AP180CTD during purification. The AP180CTD(1/3) construct (denoted as his-AP180S) was generated by introducing a stop codon in place of the codon for alanine at position 213. The plasmids for his-FUSLC (residue 1 to 163) and LAF-1 RGG (residue 1 to 168) were acquired from Addgene (https://www.addgene.org/127192/ (Fawzi laboratories) and www.addgene.org/124929/ (Hammer laboratories), respectively). The plasmid for his-FUSLC-AP180S was generated by restriction cloning the FUSLC domain into the Sal1 restriction site between the histidine tag and AP180S. The plasmid for his-AP180S-FUSLC was generated by replacing his-AP180L in the pGex4T2 vector with FUSLC-AP180S-his. First, the FUSLC-AP180S sequence from the his-FUSLC-AP180S plasmid was amplified by PCR using primers that introduced an EcoR1 cutting site to the N terminus of FUSLC-AP180S and a histidine tag, stop codon and Xho1 cutting site to the C terminus (FUSLC-AP180S-his). The his-AP180L sequence was cut out using EcoR1 and Xho1 and then EcoR1 and Xho1 digested FUSLC-AP180S-his was ligated with the remaining backbone, yielding the pGex4T2 GST-FUSLC-AP180S-his plasmid for his-AP180S-FUSLC purification. The plasmids for his-AP180L-FUSLC and his-AP180L-RGG were generated by inserting FUSLC and RGG into the C terminus of his-AP180L in a pGex4T2 vector, using restriction cloning. All constructs were confirmed by DNA sequencing.

### Protein expression and purification

His-AP180S, his-FUSLC-AP180S, his-AP180S-FUSLC, his-AP180L-FUSLC, and his-AP180L-RGG constructs were each expressed as fusion proteins with an N-terminal GST tag for increased stability. GST was subsequently removed by thrombin cleavage. All the above proteins were purified based on the following protocol. Plasmids were transformed into *E. coli* BL21Star (DE3) pLysS competent cells (NEB Cat # C2530), which were grown at 30°C to an OD 600 of 0.8. Protein expression was induced with 1 mM IPTG for 24h at 12°C, shaken at 200 rpm. The whole purification process was performed at 4ºC. The cells were pelleted from 2L cultures by centrifugation at 4,785 x g (5,000 rpm in Beckman JLA-8.1000) for 20 min. Pellets were resuspended in 100mL lysis buffer (0.5 M TrisHCl pH 8.0, 5 v/v % glycerol, 5 mM EDTA, 5 mM TCEP, 1 mM PMSF) plus EDTA free protease inhibitor tablets (1 tablet/50ml), 1.0% Triton-X100, followed by homogenization with a dounce homogenizer and sonication (5 x 2,000J) on ice. The lysate was clarified by centrifugation at 26,581 x g (18K rpm in Beckman JA-25.50) for 25 min. The clarified lysate was then applied to a 10mL bed volume Glutathione Sepharose 4B (Cytiva Cat # 17075605) column, washed with 100mL lysis buffer plus 0.2% Triton X-100, EDTA free protease inhibitor tablets (1 tablet/50ml), followed by 50mL lysis buffer. The protein was eluted with lysis buffer plus 15 mM reduced glutathione and buffer-exchanged into 50 mM TrisHCl pH 8.0, 10 mM CaCl_2_, 150 mM NaCl, 5 mM EDTA using a Zeba desalting column (Thermo Scientific cat # 89891). GST was cleaved using the Thrombin CleanCleave kit (Sigma-Aldrich cat # RECOMT) for 14 hours at 4 °C with gentle rocking. The GST-tag and any uncut protein were removed by a second Glutathione Sepharose 4B column. The resulting purified protein was concentrated using an Amicon Ultra-15 centrifugal filter (MilliporeSigma cat # UFC903024) and stored as liquid nitrogen pellets at −80 °C.

Expression and purification of his-FUSLC was carried out according to a previous report(*12*). In brief, his-FUSLC was overexpressed in *E. Coli* BL21(DE3) cells. Pellets of cells expressing his-FUSLC were harvested from 1L cultures induced with 1 mM IPTG after 4-hour incubation at 37°C and 220 RPM when OD 600 was around 0.8. The pellets were then lysed in a buffer containing 0.5 M TrisHCl pH 8, 5 mM EDTA, 5% glycerol, 10mM β-ME, 1mM PMSF, 1% Triton X-100 plus EDTA-free protease inhibitor tablet (1 tablet/50mL) for 5 min on ice and then sonicated. The cell lysates were centrifuged at 40,000 RPM for 40 min and his-FUSLC resided in the insoluble fraction after centrifugation. Therefore, the insoluble fraction was resuspended in 8M urea, 20 mM NaPi pH 7.4, 300 mM NaCl and 10 mM imidazole. The resuspended sample was then centrifuged at 40,000 RPM for 40 min. In denaturing conditions, his-FUSLC is Urea-soluble and so at this point resided in the supernatant. This supernatant was then mixed with Ni-NTA resin (G Biosciences, USA) for 1 hour at 4°C. The Ni-NTA resin was settled in a chromatography column and washed with the above solubilizing buffer. The bound proteins were eluted from the Ni-NTA resin with a buffer containing 8M urea, 20 mM NaPi pH 7.4, 300 mM NaCl and 500 mM imidazole. The purified proteins were then buffer-exchanged into 20 mM CAPS pH 11 storage buffer using 3K Amicon Ultra centrifugal filters (Millipore, USA). Small aliquots of the protein were frozen in liquid nitrogen at a protein concentration of approximately 1 mM and stored at -80 °C.

### Protein labeling

All proteins used in this study were labeled with Atto-488 or Atto-594, amine-reactive, NHS ester-functionalized fluorescent dyes. The labeling reaction took place in a 50 mM HEPES buffer at pH 7.4 for his-FUSLC and in 25 mM HEPES, 150 mM NaCl, 5 mM TCEP at pH 7.4 for AP180CTD-derived proteins. Dye was added to the protein in 2-fold stoichiometric excess and allowed to react for 30 minutes at room temperature, empirically resulting in labelling ratio from 0.8 to 1.5 dyes per protein. Labeled protein was then buffer-exchanged into 20 mM CAPS pH 11 buffer for his-FUSLC and 50 mM TrisHCl pH 8.0, 10 mM CaCl_2_, 150 mM NaCl, 5 mM EDTA, 5mM TCEP for AP180CTD-derived proteins and separated from unconjugated dye as well using 3K Amicon columns. Protein and dye concentrations were monitored using UV–Vis spectroscopy. Labeled proteins were dispensed into small aliquots, flash frozen in liquid nitrogen and stored at -80°C. For all experiments involving labeled protein, a mix of 90% unlabeled / 10% labeled protein was used.

### GUV preparation

GUVs were made of 83% POPC, 15% Ni-NTA, 2 mol% DP-EG10 biotin. An additional 0.1 mol% Texas Red DHPE lipids were added for visualization if needed. GUVs were prepared by electroformation according to published protocols(*41*). Briefly, lipid mixtures dissolved in chloroform were spread into a film on indium-tin-oxide (ITO) coated glass slides (resistance ∼8-12 W sq-1) and further dried in a vacuum desiccator for at least 2 hours to remove all of the solvent. Electroformation was performed at 55°C in glucose solution. Glucose solutions with different osmolarity were used to match the osmolarity of buffers with different NaCl concentrations. To be specific, 560 milliosmole glucose solution was employed for making GUVs used in 250 mM and lower NaCl buffers. 940 milliosmole glucose solution was used for GUVs in 500 mM NaCl buffer. For GUVs used under 1 M NaCl buffer and 250 mM NaCl plus 1.5 M Urea, 1800 milliosmole glucose solution was adopted to adapt to the high osmotic pressure. The voltage was increased every 3 min from 50 to 1400 mV peak to peak for the first 30 min at a frequency of 10 Hz. The voltage was then held at 1400 mV peak to peak for 120 min and finally was increased to 2200 mV peak to peak for the last 30 min during which the frequency was adjusted to 5 Hz. GUVs were stored in 4°C and used within 3 days after electroformation.

### GUV tethering and sample preparation

GUVs were tethered to glass coverslips for imaging as previously described(*42*). Briefly, glass cover slips were passivated with a layer of biotinylated PLL-PEG, using 5 kDa PEG chains. GUVs doped with 2 mol% DP-EG10-biotin were then tethered to the passivated surface using neutravidin.

PLL-PEG was synthesized by combining amine reactive PEG and PEG-biotin in molar ratios of 98% and 2%, respectively. This PEG mixture was added to a 20 mg/mL mixture of PLL in a buffer consisting of 50 mM sodium tetraborate (pH 8.5), such that the molar ratio of lysine subunits to PEG was 5:1. The mixture was continuously stirred at room temperature for 6 h and then buffer exchanged into 25 mM HEPES, 150 mM NaCl (pH 7.4) using ZebaTM Spin Desalting Column (ThermoFisher Scientific).

Prior to tethering, the osmolality of the GUV solution and experimental buffers were measured using a vapor pressure osmometer (Wescor). Buffer used for dilution and rinsing was 25 mM HEPES pH 7.4 with corresponding NaCl concentrations. Osmolarity balance was maintained by the addition of glucose to the buffer.

Imaging wells consisted of 5 mm diameter holes in 0.8 mm thick silicone gaskets (Grace Bio-Labs). Gaskets were placed directly onto no.1.5 glass coverslips (VWR International), creating a temporary water-proof seal. Prior to well assembly, gaskets and cover slips were cleaned in 2% v/v Hellmanex III (Hellma Analytics) solution, rinsed thoroughly with water, and dried under a nitrogen stream. In each dry imaging well, 20 μL of PLL-PEG was added. After 20 min of incubation, wells were serially rinsed with appropriate buffer by gently pipetting until a 15,000-fold dilution was achieved. Next, 4 μg of neutravidin dissolved in 25 mM HEPES, 150 mM NaCl (pH 7.4) was added to each sample well and allowed to incubate for 10 minutes. Wells were then rinsed with the appropriate buffer to remove excess neutravidin. GUVs were diluted in appropriate buffer at ratio of 1:13 and then 20 μL of diluted GUVs was added to the well and allowed to incubate for 10 minutes. Excess GUVs were then rinsed from the well using the appropriate buffer and the sample was subsequently imaged using confocal fluorescence microscopy.

### GUV fluorescence imaging

Imaging experiments were performed using a spinning disc confocal super resolution microscope (SpinSR10, Olympus, USA) equipped with a 1.49 NA/100X oil immersion objective. Laser wavelengths of 488 and 561 nm were used for excitation. Image stacks taken at fixed distances perpendicular to the membrane plane (0.5 μm steps) were acquired immediately after GUV tethering and again after protein addition. Images taken under deconvolution mode were processed by the built-in deconvolution function in Olympus CellSens software (Dimension 3.2, Build 23706). At least 30 fields of views were randomly selected for each sample for further analysis prior to and after the addition of protein, respectively. Imaging was proceeded 5 min after adding proteins to achieve protein binding and reaching a relatively equilibrium state.

### Statistical analysis

All GUV experiments were repeated 3 times for each condition reported. ImageJ was employed to analyze confocal images. At least 100 GUVs were examined under each condition. The diameter of each inward tubule was determined by drawing a line perpendicular to the tubule at three different places along its length and calculating the average diameter. To assess the significance of comparisons between conditions, an unpaired t-test was performed. Error bars in graphs represent either standard error or standard deviation as stated in figure captions.

## SUPPLEMENTARY INFORMATION

### Supplementary Figures

**Figure S1.**
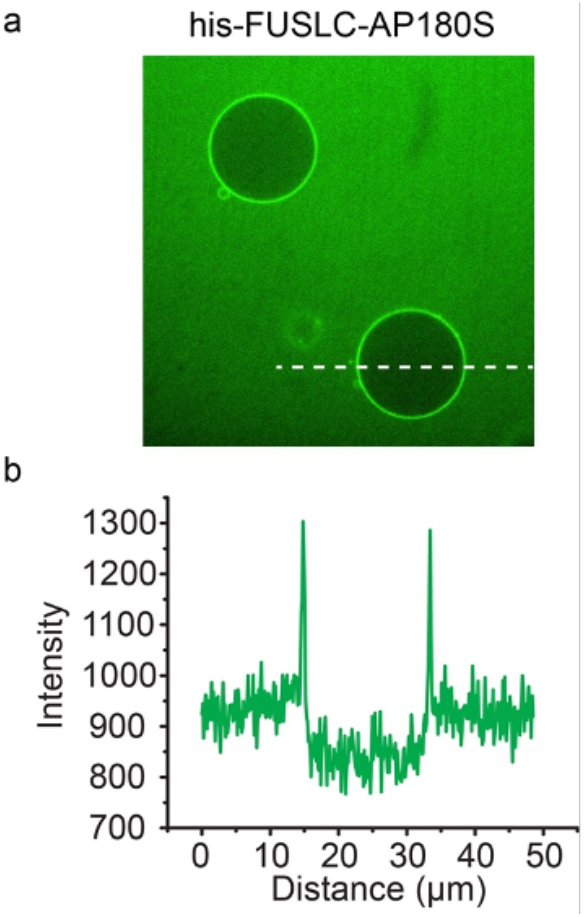
(a) Representative confocal images showing protein exclusion from the inner lumen of GUVs. (b) Intensity distribution of protein channel along the dashed line shown in the image.

**Figure S2.**
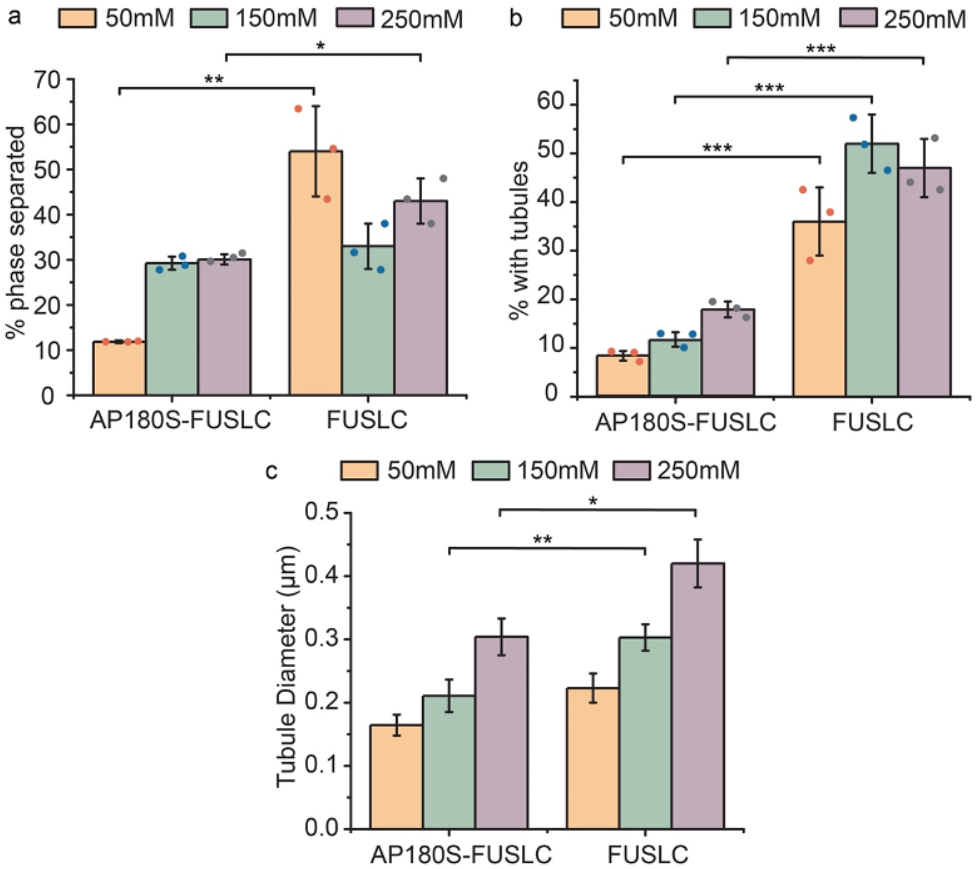
Bar chart comparison of percentage of GUVs displaying protein phase separation (a), inward tubules (b) and average inward tubule diameter (c) when incubated with his-AP180S-FUSLC and his-FUSLC under different salt concentrations. The data for his-FUSLC are cited from our previous report^12^. Error bars in (a) and (b) represent the standard deviation of three independent trials (indicated by the dots). Error bars in (c) correspond to standard error of all data points measured under each condition. Statistical significance was tested using an unpaired, two-tailed student’s t test. *: P < 0.05, **: P < 0.01, ***: P < 0.001. GUV composition is 83 mol% POPC, 15 mol% DGS-NTA-Ni, 2 mol% DP-EG10 biotin, and 0.1 mol% Texas Red-DHPE.

**Figure S3.**
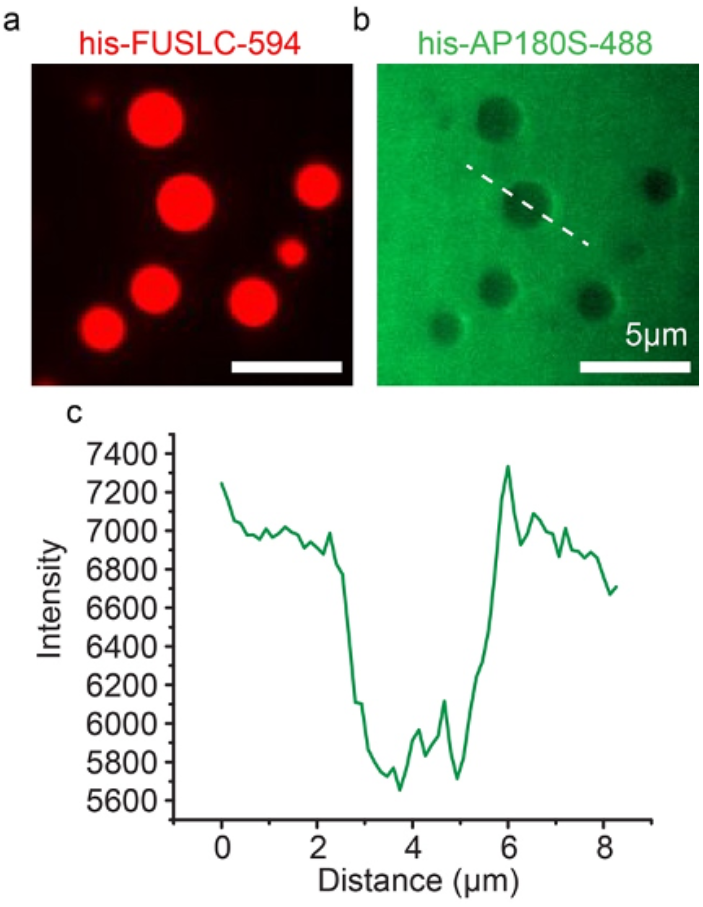
(a, b) Representative image of AP180S exclusion from his-FUSLC droplets in solution. (c) Intensity distribution of his-AP180S along the white dashed line across the his-FUSLC droplet in panel b. his-FUSLC concentration is 25 μM and his-AP180S is 5 μM. Experiment was done in 25mM HEPES, 150mM NaCl, pH 7.4 buffer.

**Figure S4.**
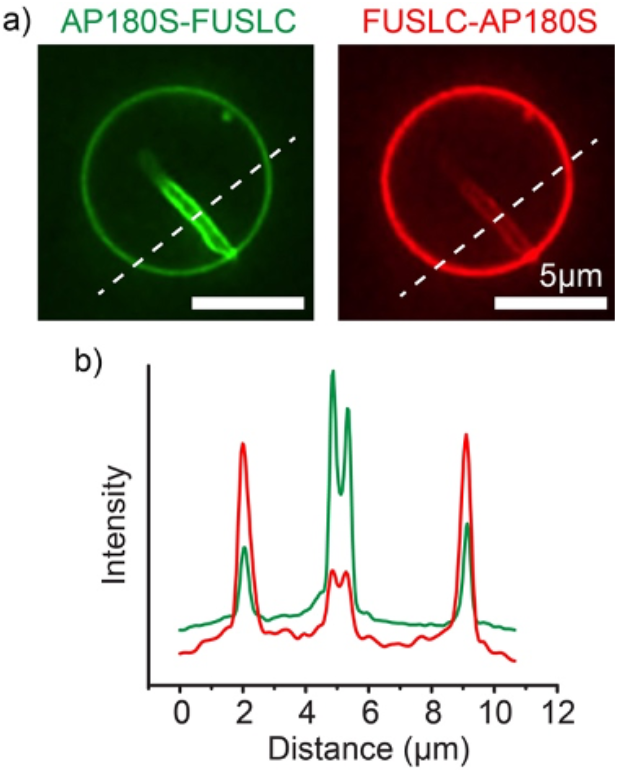
AP180S-FUSLC is more enriched in the inward protein-lined tubules than FUSLC-AP180S. (a) Representative super-resolution images of protein-lined tubules when GUVs were incubated with 1 μM of his-AP180S-FUSLC and 0.1 μM of his-FUSLC-AP180S at the same time. (b) Intensity distribution along the dashed line across the GUV.

### Supplementary Information about Simulations

To the first order, the total system energy can be written as:

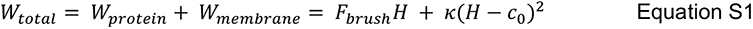

where *F*_*brush*_ is the protein energy density, *H* is the mean curvature of the membrane, κ is the bending modulus, and *c*_0_ the spontaneous curvature. The first term, the product of protein energy density and mean curvature, captures the moment arm generated by the protein steric interactions which are offset from the membrane surface. The second term is the conventional Helfrich-Canham-Evans hamiltonian which captures the membrane’s bending elasticity.(*25-27*)

The physics of interacting polymers grafted on surfaces has been explored in detail and the brush polymer theory has been developed to model the contributions to the polymer energy density (*21-24*). Following the theory, the protein energy density, F brush, is defined as,

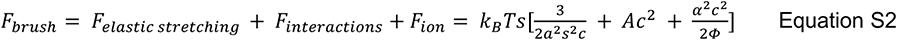

where *s* is the local area per protein, *a* is the Kuhn length, *c* the protein monomer concentration (considered to be an amino acid residue for the IDPs of interest), *A* the second virial coefficient, *Φ* the bulk ionic concentration, *k*_*B*_*T* is the Boltzmann constant and temperature, and *α* the degree of ionization. While a freely jointed polymer chain is expected to form a glob like structure which maximizes its configurational entropy(*43*), high density packing conveyed by the area occupancy and protein concentration, can lead to stretching and subsequent entropy loss which is captured by the first term. The free energy density of the short-ranged interactions which can drive phenomena such as steric pressure and condensation is modeled by the virial expansion truncated to second order, the second term of Eq. S2. Here we note that *A*, the second virial coefficient, is a complex function of the specific protein chemistry/identity(*44, 45*) and can take a positive or negative value corresponding to repulsive or attractive behavior respectively. Conventional biochemistry and statistical mechanics further imply that *A* is a function of the local chemical environment which can screen the strength of interactions. In lieu of detailed experimental characterization of *A* given the combinatorial conditions, we will instead systematically vary the value of *A* across a range of values corresponding to net protein attraction/aggregation and repulsion. The third term of Eq. S2, captures the loss of entropy of ions partitioning into the brush layer as a result of Donnan’s equilibrium. From the brush polymer theories, this term corresponds to the so-called quenched or strongly dissociating condition which assumes that the polymer has a fixed ionization extent given by *α*.

To interrogate how the IDP layer and membrane couple to drive spontaneous curvature we can study how the energy of the system changes with respect to changing mean curvature, ∂*W*/∂*H*.

The stationary point where 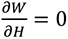, corresponds to the minima and is given by

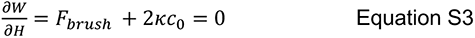

Rearranging, we obtain the classic relationship linking the moment of the protein interactions with the spontaneous curvature and bending rigidity,

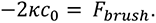

For a quantitative evaluation of this relationship, we further assume and prescribe values for the parameters. The Kuhn length, *a*, is twice the persistence length of a polymer. For an IDP we approximate this as twice the length of a residue, ∼1 nm.(*46*) To relate the end-to-end stretch distance of the grafted IDP (i.e., thickness of the brush layer), *d*, to polymer area occupancy we write a conservation equation,

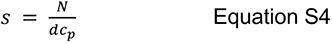

where *N* is the number of amino acids, and *c*_*p*_ is the protein monomer concentration. We further assume that the degree of ionization for each protein is around 10%, α = 0.1, the brush layer is 20 nm thick, and prescribe 150 mM ion concentration. Under these conditions we vary the second virial coefficient, *A*, from -10 – 10 nm^3^ corresponding to net attractive (e.g., FUSLC) and repulsive (e.g., AP180S) conditions respectively. We find that the extent of spontaneous curvature induced is a function of protein concentration in line with the experimental observations, Figure 6b.

To evaluate how the spontaneous curvature influences the geometry of a membrane tube, we followed the approach outlined by Shurer, Derenyi and colleagues(*9, 47*). Considering the special case of a cylindrical membrane tube experiencing a pulling force at zero osmotic pressure, the Helfrich-Canham-Evans free energy is given by,

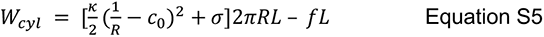

where κ is the bending modulus, *R* is the radius of the cylinder, *c*_0_ is the spontaneous curvature, σ is the tension, *f* is the pulling force acting over length *L*. At equilibrium, where 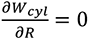 and ∂*W*_cyl_/∂*L =* 0, the tube radius is

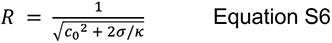

and the pulling force,

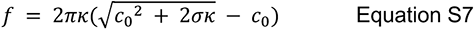

Assuming a canonical bending rigidity value of 20 k_B_T and membrane tension 0.01 mNm^-1^, and substituting the predicted values from Figure 6b, we obtain the radius and force predictions in Figure 6c and Figure S5.

For a proof of principle of inward and outward tube formation with pearling, we used Mem3DG(*28*) to build a model in three dimensions without assumptions of axisymmetry. Mem3DG is a framework and simulation engine that enables us to solve the governing equations of membrane bending. Using principles from discrete differential geometry, we compute the energy of a geometric configuration and vertexwise forces in a consistent manner with traditional physics approaches; coupled with an energy minimization or time integration scheme, we get from the model a trajectory of the evolving domain subject to the bending and other prescribed physics. For the demonstration, we start from a spherical vesicle and apply a weak Gaussian point force, decaying in time, in an inward or outward direction to induce initial tube formation. Proteins which impart spontaneous curvature matching the direction of the tube bind and support the extrusion of pearls. Driven by the membrane tension and protein spontaneous curvature, the resulting configurations, shown in Figure 6d, exhibit pearled tubules similar to those observed in experiments, (Figure 2c, 4c) suggesting that our simple physical model captures the main features of our experimental system.

All parameters for the Mem3DG and simple mechanical models are archived on GitHub: https://github.com/RangamaniLabUCSD/2023-IDP-bending. Mem3DG source code corresponding to commit 361affa9423d44f3cf239585ac350340a212b1f8 used to run the model can also be obtained from GitHub https://github.com/RangamaniLabUCSD/Mem3DG/.

**Figure S5.**
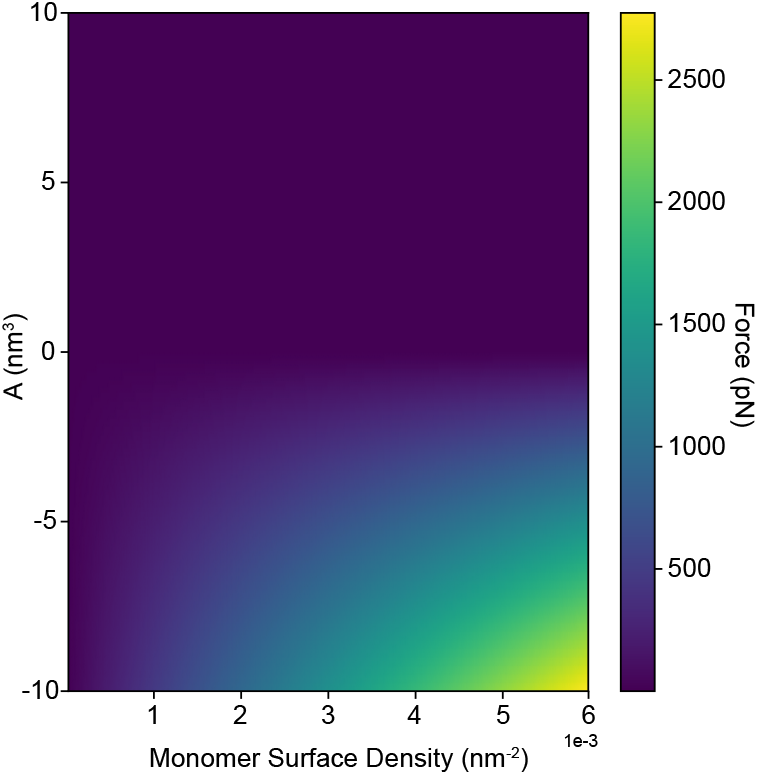
The equilibrium pulling force required to sustain a tube coated by differing surface concentrations of protein. The vertical axis is the second virial coefficient, A, which represents the net attraction—repulsion of the protein.

